# Characterizing Clinical Toxicity in Cancer Combination Therapies

**DOI:** 10.1101/2025.04.13.648641

**Authors:** Alexandra M. Wong, Lorin Crawford

## Abstract

Predicting synergistic cancer drug combinations through computational methods offers a scalable approach to creating therapies that are more effective and less toxic. However, most algorithms focus solely on synergy without considering toxicity when selecting optimal drug combinations. In the absence of combinatorial toxicity assays, a few models use toxicity penalties to balance high synergy with lower toxicity. However, these penalties have not been explicitly validated against known drug-drug interactions. In this study, we examine whether synergy scores and toxicity metrics correlate with known adverse drug interactions. While some metrics show trends with toxicity levels, our results reveal significant limitations in using them as penalties. These findings highlight the challenges of incorporating toxicity into synergy prediction frameworks and suggest that advancing the field requires more comprehensive combination toxicity data.

## 1 Introduction

Cancer is a significant cause of mortality both in the United States as well as globally [1], and its incidence is expected to increase over the next fifty years [2]. To address drug resistance, the cause of over 90% of cancer deaths [3, 4], scientists have successfully developed combination therapies to combat tumor heterogeneity and narrow possible mechanisms for therapeutic escape [5, 6]. However, discovering novel combination therapies involves costly high-throughput screening assays, which become increasingly more expensive as the number of potential drug candidates grows [7, 8]. To mediate this combinatorial explosion, *in silico* approaches have taken advantage of the large-scale multi-omic and cancer drug databases [9–12] to learn to predict new therapeutic combinations.

Many of these *in silico* methods use machine learning to predict a “synergy score” [13–22], which is a quantitative measure meant to optimize for drug-drug interactions (DDI) that produce a combined effect greater than the “additive” effect (i.e., the sum of their individual effects). However, there is increasing evidence that synergy scores may not be the best metric to optimize over. First, additivity alone has been shown to be predictive of the efficacy of many approved cancer drug combinations [23, 24]. Second, cancer combination therapies can better accommodate the diverse nature of patient populations, increasing the likelihood that at least one drug will be effective for each patient, rather than relying on a synergistic effect in any single individual [24]. Additionally, what can be considered “synergistic” is variable. For example, the DrugComb database reports seven different synergy scoring methods [9], reflecting that defining synergy remains an open question. Thus, optimizing for synergy scores alone may not be the best primary focus for predictive computational models.

Another significant hurdle in cancer therapeutics development is toxicity, which is the second biggest reason for clinical trial failure [25–27]. Even after a therapy is approved, intolerable side effects can contribute to the early discontinuation of a patient’s treatment schedule [28–30]. Toxicity plays a critical role in clinical translation, and drug synergy is thought to reduce toxicity based on the principle that higher synergism allows for lower dosages of each drug to create the same efficacy on the cancer, reducing the potential side effects from either individual drug [6, 31]. While many methods cite reducing toxicity as a motivation for finding synergistic drug combinations, most computational models only focus on predicting a synergy value without regard to adverse drug effects [13, 14, 16–19, 32]. Fewer even attempt to integrate toxicity. One approach, SynToxProfiler, studied a prioritization scheme to balance synergy with toxicity and efficacy when ranking optimal combination candidates [33]. Notably, some of their top-performing combinations had negative synergy scores (i.e., combinations that would have been missed when selecting by synergy scores alone). SynToxProfiler created toxicity scores through experimental measurements of drug combinations tested on healthy control cells. However, this type of combinatorial toxicity data is not often available for many of the large cancer drug combination datasets that are widely used in the field.

Absent that data, some combination prediction methods have sought to incorporate toxicity information in other ways. Chen et al. [21] and Hosseini and Zhou [34] used clinical side effects of drugs from the SIDER database as drug features in their combination prediction, but they did not explicitly attempt to minimize toxicity. These approaches also did not consider known adverse effects from the drug *combination*, as SIDER only contained adverse effects for single drugs. Other methods added toxicity penalties in their loss functions when predicting synergistic combinations, seeking to balance high synergy with low toxicity. DeepTraSynergy by Rafiei et al. [15] and GraphSynergy by Yang et al. [35] created novel loss functions based on the idea that greater drug target and pathway overlap leads to higher toxicity [36]. DeepTraSynergy also incorporated drug structure representations into its toxicity penalty because similarity of drug structures is also thought to increase the chances of adverse DDIs [37]. However, whether these metrics correspond to clinical toxicity remains uncharacterized.

In this study, we explore how well synergy scores correlate with clinically known combination toxicity. We ask three key questions. First, is there a relationship between synergy scores and toxicity levels of DDIs? This question is critical for understanding if current methods may be optimizing for more or less toxic combinations when prioritizing only by the synergy score. Second, can overlap of drug targets and pathways explain toxicity and synergy? This assessment seeks to confirm how robust target overlap is as a measure of toxicity. Finally, do common methods used in toxicity penalties correlate with known DDIs? This analysis investigates whether toxicity penalties are sufficient for balancing efficacy and safety in combination candidates. In our analysis, we show that while current measures may correlate with general toxicity trends, they are insufficient in capturing the complexity of clinically known adverse DDIs and that there is a need for more explicit combinatorial toxicity data.

## 2 Methods

We used the DrugBank [38] and DDInter [39] databases to retrieve toxicity level information for drug combinations. Our study has three major parts: (i) an analysis of the relationship between synergy scores and toxicity, (ii) an analysis of the biological mechanisms underpinning toxicity and synergy, and (iii) an analysis of whether toxicity penalty methods correlate with known toxicity (see schematic overview in Figure 1). Detailed descriptions of the databases, scoring methods, and statistical tests used in our analyses are provided in the sections below.

**Figure 1:**
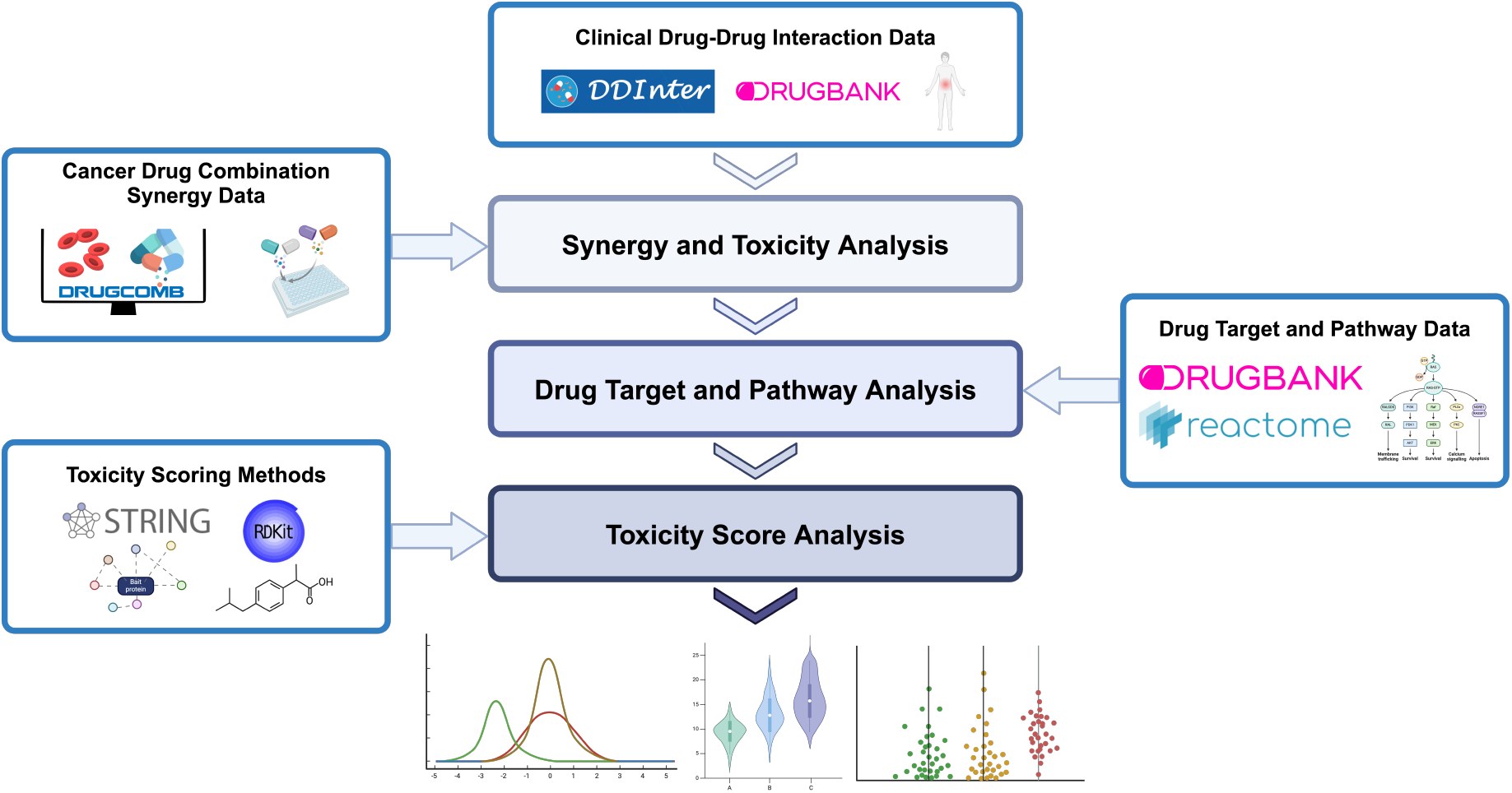
Overview of the databases and analyses conducted in this study. We first retrieved the drug-drug interactions (DDI) toxicity data from the DrugBank and DDInter databases. For the synergy and toxicity analysis, we used the toxicity data in combination with the synergy scores from the DrugComb [9] database. For the drug target and pathway analysis, we retrieved drug target data from DrugBank and pathway information from Reactome [40]. For the toxicity score analysis, we used STRING [41] to create drug target distance metrics and acquired drug structure representations through RDKit [42]. Illustration created by AMW in BioRender [43].

### 2.1 Clinical drug-drug interaction data

We used two different databases containing information about known DDI data. Information on all database versions and where to access them can be found in the section below entitled “Data and Code Availability”. Our primary source of toxicity information was DrugBank, where we retrieved over 1.4 million DDIs under an academic license [38]. These data included the name of each drug in the drug pair, the drug targets for each drug, and the severity level of the DDI (encoded as either 0, 1, or 2 depending on whether the severity was minor, moderate, or major, respectively). We chose DrugBank as our primary dataset due to its more comprehensive size and frequent updates. To assess the reproducibility of our results, we also conducted the same set of analyses using DDInter, which contains 160,235 DDIs [39]. DDInter represented its DDIs by mapping drug pairs to one out of three severity levels (minor, moderate, and major). For consistency and ease of comparison between databases, we converted DrugBank’s 0, 1, and 2 severity values to minor, moderate, and major.

### 2.2 Cancer drug combination synergy data

To evaluate the relationship between synergy scores and the severity of toxicity, we used the cancer drug combination synergy scores in DrugComb [9]. We chose DrugComb for its unification of several pairwise drug combination datasets [10, 11, 44, 45] and synergy scoring methodologies. All synergy scores in DrugComb were calculated by assessing the degree of deviation of an observed response from the expected effect of the “additive” interaction. This meant that a synergy score of less than zero is antagonistic, equal to zero is additive, and greater than zero is synergistic. DrugComb reported five distinct models of synergy scores that formulate additivity differently: Bliss, Highest Single Agent (HSA), Loewe, S, and Zero Interaction Potency (ZIP). The S synergy score model has three variations in how it was calculated: S_max, S_mean, or S_sum. Our study included all seven synergy scores. We provide the formulas for each synergy score below, which can also be found on the DrugComb documentation [9, 46, 47].

Let *S*_*model*_ represent the synergy score as calculated by a model (i.e., one of Bliss, HSA, Loewe, ZIP, max, mean, or sum). Let D be the set of drugs and *R*_*A*_ be the measured response of individual drug *A* ∈ *𝒟*. We defined *E*_*AB*_ as the measured combination effect between drugs {*A, B*} ∈ *𝒟*.

#### Bliss

The Bliss Independence Model assumes that drugs act independently through different mechanisms. Namely,

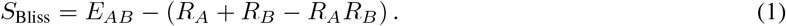

Although the Bliss synergy score is summed over dose-response relationships, it offers an intuitive interpretation of what is considered ‘greater than additive’ activity.

#### HAS

The Highest Single Agent (HSA) model is the most simplistic synergy score. HSA calculates the difference between combination effects and the most effective single drug

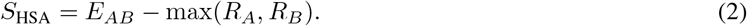

While straightforward to implement and interpret, HSA is more permissive in identifying synergy and also does not consider dose-response curves, potentially overestimating synergistic effects in many scenarios.

#### Loewe

The Loewe Additivity Model assumes that the combined effect of two drugs can be predicted if the doses of each drug are adjusted to account for their relative potencies

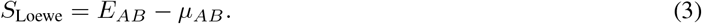

Let concentrations of drugs *A* and drug *B* be denoted by *C*_*A*_ and *C*_*B*_, respectively. The expected effect *µ*_*AB*_ for a given drug combination *AB* must satisfy the following

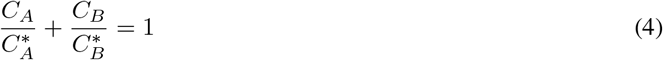

where 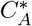 and 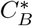 are the doses of drug *A* and *B* that produce the same effect individually as they do in combination. To better understand 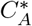 and 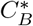, we can substitute them by using dose-response curves described by four parameter log-logistic curves. The condition that the drug combination must satisfy then becomes the following

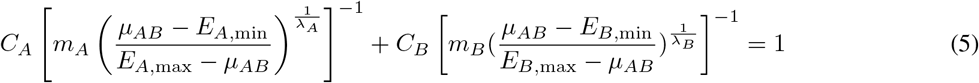

where *m*_*A*_ and *m*_*B*_ are the half-maximal inhibitory concentrations for drugs *A* and *B* respectively, the values {*E*_*A*,min_, *E*_*A*,max_} ∈ [0, 1] are the minimal and maximal effects of the drug, and *λ*_*A*_ and *λ*_*B*_ are the sigmoidicity shape parameters of the dose response curves. Unlike the previous scores, the Loewe synergy score incorporates dose-response relationships of individual drugs, but is more computationally intensive.

#### ZIP

The Zero Interaction Potency (ZIP) score combines the Bliss and Loewe models by calculating the expected combination effect of two drugs under the assumption that they do not potentiate each other. Namely,

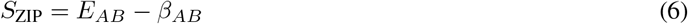

where the expected effect *β*_*AB*_ for a drug combination *AB* is given by the following

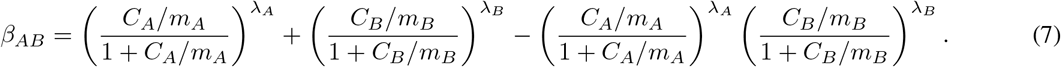

This unified approach addresses the limitations of both parent models but is more complicated to interpret. The ZIP score also requires comprehensive dose-response matrices.

#### S Synergy Scores

The S scores are based off of a Combination Sensitivity Score (CSS) that integrates dose response curves to determine the sensitivity of a drug combination. The S scores are calculated as deviations of the observed drug combination effect from the expected effect if the drugs do not interact. Namely,

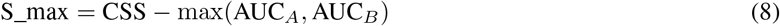

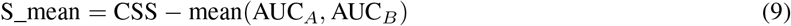

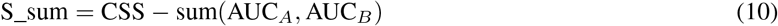

where AUC_*A*_ and AUC_*B*_ are areas under the monotherapy dose-response curves. Each S score uses a different reference model: S_max uses the effect of the most potent single drug (similar to HSA), S_mean uses the average effect of individual drugs, and S_sum uses the sum of individual effects.

### 2.3 Data preprocessing

We used the following procedures to preprocess the data used in this study. First, we focused our study on cancer drug combinations from DrugComb where all seven synergy scores were available. For reproducibility, we then filtered the DrugComb combinations to create two separate datasets: one for the combinations with known toxicity levels in DrugBank, another for DDInter. For both datasets, we then also limited the combinations to drugs that had known drug target information (via DrugBank) and known UniProt IDs for mapping to the Reactome pathways and STRING protein-protein interaction network (PPIN). When all filtering was finished using the DrugBank toxicity levels, there were 526 unique drugs, 53,516 unique (drug *A*, drug *B*, cell line *C*) triplicates invariant to order of (drug *A*, drug *B*) versus (drug *B*, drug *A*), and 186 unique cell lines. Using the DDInter toxicity data, the end result of preprocessing left 331 drugs, 23,415 unique drug-drug-cell line triplicates, and 149 cell lines.

### 2.4 Synergy scores and toxicity analysis

In the first analysis, we examined whether synergy scores have any relationship with DDI severity. Using the final processed datasets for both DrugBank and DDInter, we first characterized the percentages of toxicity categories among synergistic combinations (Figure 2A and 2C). We then tested whether drug combination synergy scores were normally distributed using scipy’s `normaltest` function [48], based on the D’Agostino and Pearson’s test [49, 50]. Each of the seven synergy scores did not appear to follow Gaussian distributions (*p* ≤ 0.05 where the null hypothesis assumes a sample is drawn from a normal), leading us to use non-parametric tests. We then split the synergy scores by minor, moderate, and major toxicity levels as determined by DrugBank or DDInter. Using the Kruskal-Wallis test (via `scipy.stats.kruskal`) [48, 51], we first asked if toxicity categories had significant differences in the medians of their synergy score distributions. For post-hoc analysis, we applied Dunn’s test [52, 53] with Bonferroni correction using `scikit_posthocs.posthoc_dunn` [54] to assess whether there were significant differences in the medians between pairwise categories (i.e., between major and moderate, moderate and minor, or major and minor). To interrogate whether optimizing for synergy had any bearing on lowering toxicity, we used the Jonckheere-Terpstra test [55, 56] to assess the directionality of trends between the toxicity levels and synergy scores. In other words, we tested if optimizing for higher synergy scores was correlated with lower toxicity. First, we tested if the medians of the synergy score distributions increase as toxicity levels decrease. We then computed whether the medians of synergy scores increase as toxicity levels increase. All statistical tests and *p*-values are present in Supplementary Table 1. Box plots for the synergy scores broken down by toxicity level are available in Figure 2 and Supplementary Figures 1 and 2.

**Figure 2:**
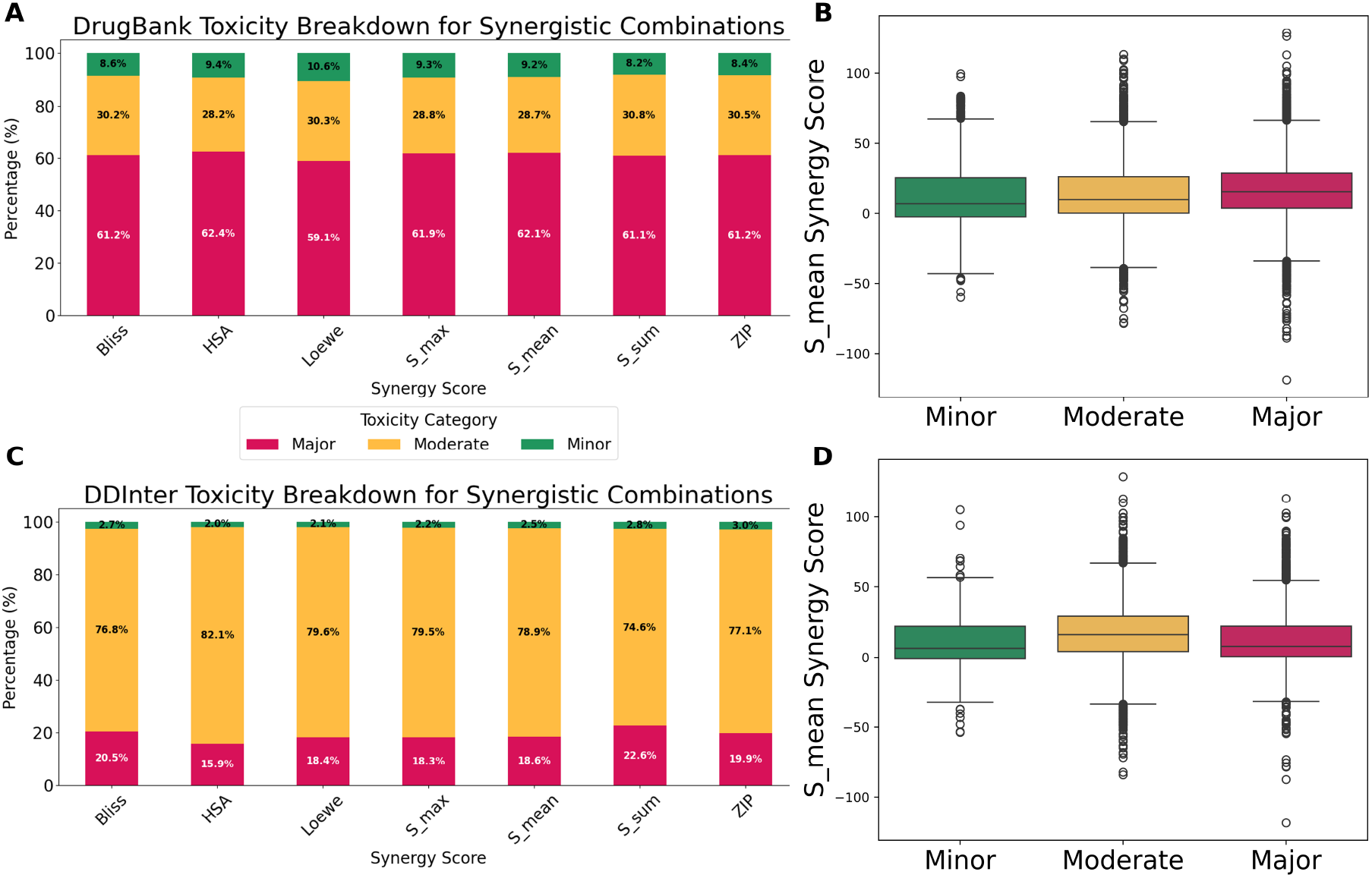
Synergistic drug combinations trend with higher toxicity in DrugBank but not DDInter. Panel **(A)** shows the percentage breakdown of toxicity categories for synergistic combinations in DrugBank. Panel **(B)** displays boxplots showing the distributions of S_mean synergy score from the DrugBank dataset when split across toxicity categories: the median of each distribution is denoted by the black horizontal line and the interquartile range is bounded by the box. Panel **(C)** shows the percentage breakdown of toxicity categories for synergistic combinations in DDInter. Panel **(D)** displays boxplots showing the distributions of S_mean synergy score from the DDInter dataset when split across toxicity categories. Additional DrugBank synergy score distributions are presented in Supplementary Figure 1 and DDInter synergy score distributions are presented in Supplementary Figure 2.

### 2.5 Drug target and pathway analysis

In the second component of our analysis, we investigated the mechanisms that underpin toxicity and synergy. We used drug targets and their UniProtIDs from DrugBank [38]. We then mapped the targets to the pathways they belong to in the Reactome database [40]. We limited our search of Reactome pathways to those in *Homo sapiens*. Reactome stores a hierarchical structure of pathways, so we included two categories of pathway levels when computing the overlap of pathways between two drug targets. The “lowest pathway level” is the set of pathways in Reactome that belongs to the lowest level of the hierarchy (i.e., the bottom child nodes in the Reactome pathway tree). The “all pathway levels” category includes all pathways in Reactome, regardless of where in the hierarchy it belongs. Drug targets, their pathways at the lowest level, and their pathways at all levels were used in the drug target and pathway analysis section.

We first examined whether drug target and pathway overlap was correlated with DDI severity levels. For each dataset (DrugBank and DDInter), we computed the Jaccard Similarity, a metric of overlap [57, 58], for all drug combinations for each of these three categories. For example, for each (drug *A*, drug *B*, cell line *C*) triplicate, we find the set of all targets for drug *A, T*_*A*_, the set of all targets for drug *B, T*_*B*_, and compute the Jaccard Similarity *J* (*A, B*) between the two sets. The Jaccard Similarity is calculated as

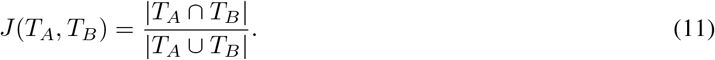

This process is also applied to the drug targets’ Reactome pathways, resulting in Jaccard Similarity scores for drug targets, lowest Reactome pathways, and all Reactome pathways. We then tested each of these similarity scores for normality using the D’Agostino and Pearson’s test. Both DrugBank and DDinter were not normal (*p* ≤ 0.05), so we used the Kruskal-Wallis test to assess if toxicity categories had significant differences in the medians of the drug target or pathway Jaccard Similarities. We performed post-hoc analysis via Dunn’s test with Bonferroni correction. To determine whether there was a significant ordering of Jaccard Similarity medians when increasing or decreasing the toxicity level, we used the Jonckheere-Terpstra test. All statistical test outcomes can be found in Supplementary Table 2. A schematic depicting the steps for this analysis is present in Figure 3A. Strip plots to show the Jaccard Similarity distributions are shown in Figure 3B-3G.

**Figure 3:**
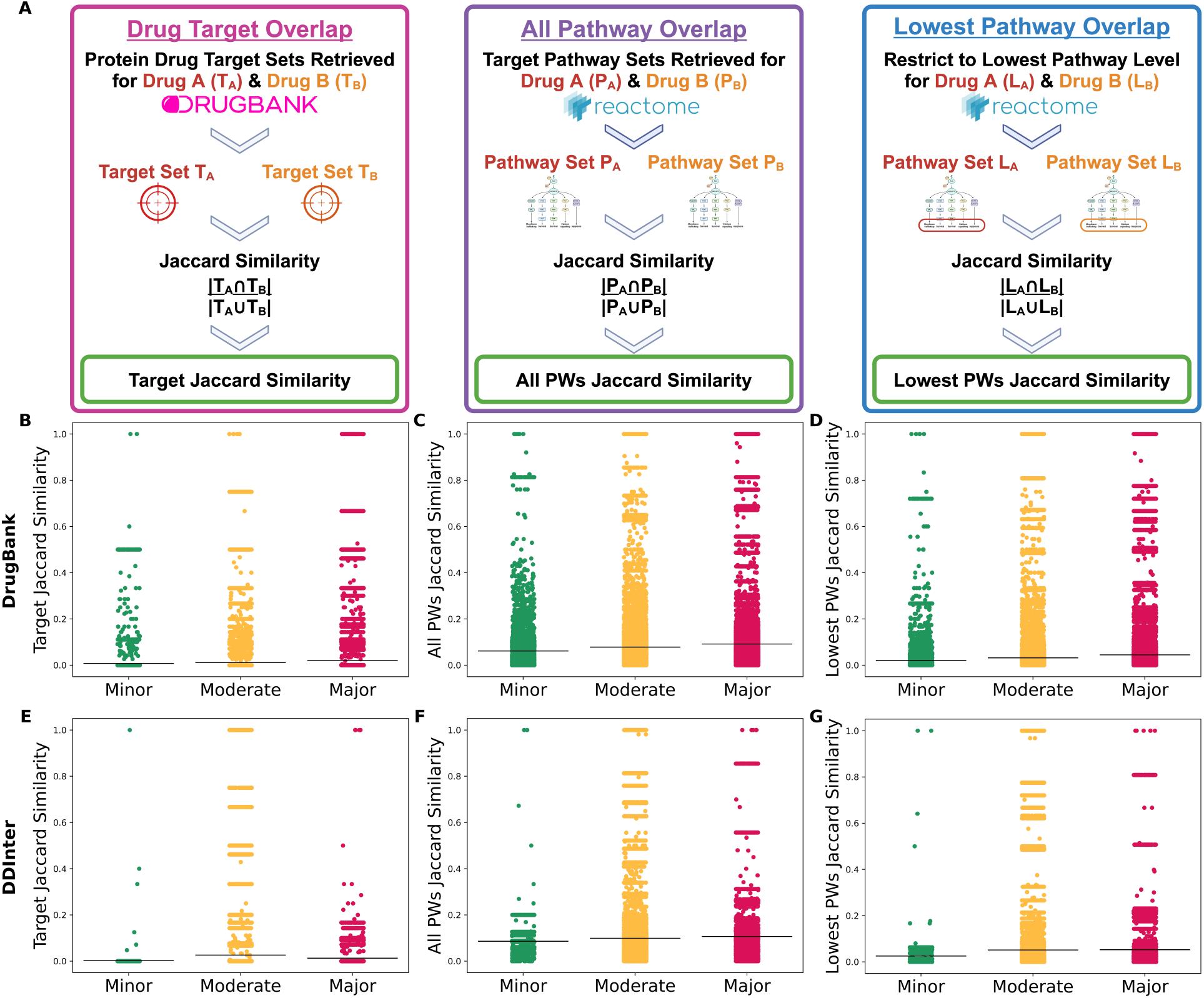
Drug target and pathway overlap metrics do not consistently separate toxicity categories. Panel **(A)** shows a flow chart illustrating how each of the Target, All Pathways (All PWs), and Lowest Pathways (Lowest PWs) Jaccard Similarities were calculated. The remaining panels are strip plots displaying the distributions of drug combination target and target pathway overlap when split across toxicity categories. The mean of each distribution is denoted by the black horizontal line. All distributions representing the Minor toxicity are in green, the Moderate in yellow, and the Major in red. The top row including panels **(B), (C)**, and **(D)** used DrugBank toxicity categories, while the bottom row of panels **(E), (F)**, and **(G)** correspond to DDInter. Panels **(B)** and **(E)** show the Jaccard distributions of toxicity categories based on the set of target proteins for each drug in a drug pair. Panels **(C)** and **(F)** display the Jaccard Similarity distributions based on all Reactome pathways, while Panels **(D)** and **(G)** restrict Reactome to its lowest level pathways.

We also examined whether there was a correlation between drug target and pathway overlap with synergy scores. We used the same Jaccard Similarity metrics for drug targets, lowest Reactome pathways, and all pathways to compute a Pearson [59] and Spearman [60] correlation with each of the synergy scoring methods. We then created scatter plots with best fit lines and their R-squared (*R*^2^) [61] value. We completed these steps both for the preprocessed DrugBank and DDInter datasets, which can be found in Figure 4, Supplementary Table 3, and Supplementary Figures 3 and 4.

**Figure 4:**
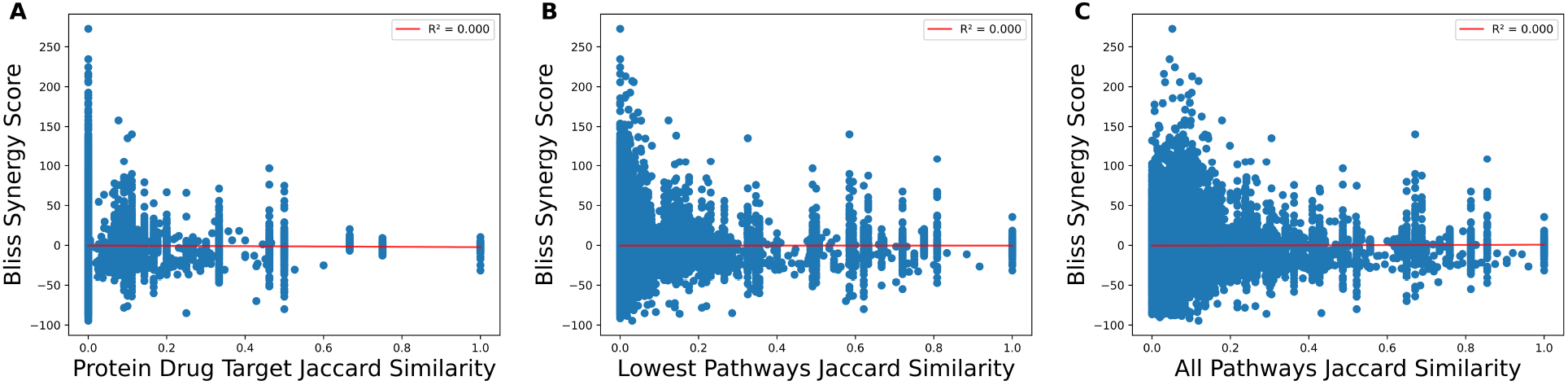
Synergy scores show no correlation with drug target overlap metrics from DrugBank. Scatter plots showing the Bliss synergy score versus three drug target overlap metrics for combinations present in both DrugComb and DrugBank. Panel **(A)** corresponds to the Jaccard Similarity of drug combinations’ protein targets. Panel **(B)** shows the Jaccard Similarity of drug target pathways when restricted to the lowest level of Reactome. Panel **(C)** displays the Jaccard Similarity of drug target pathways at all levels of Reactome. Each plot also contains a red line for the best fit line, with the *R*^2^ present in the legend, which are all zero. See Supplementary Figure 3 for all other synergy scores.

### 2.6 Toxicity score analysis

Some cancer drug combination prediction models integrate toxicity scores into their loss functions to balance optimizing for high synergy while minimizing toxicity. We assessed three metrics to test key components of toxicity scores (Figure 5A). First, some methods use drug structure similarity as a toxicity penalty term [15]. To assess this, we retrieved the SMILES [62] string representations for all drug structures from DrugBank. We then used RDKit [42] to map the SMILES representations to Morgan Fingerprint 2048-bit vectors [63]. For each drug combination (say drug *A* and drug *B*), we used the corresponding Morgan Fingerprint vectors (*M*_*A*_, *M*_*B*_), and calculated the Tanimoto Similarity *T* (*M*_*A*_, *M*_*B*_) [64]. The Tanimoto Similarity is equivalent to the Jaccard Similarity. For bit vectors like Morgan Fingerprint representations, *T* (*M*_*A*_, *M*_*B*_) can be expressed as

**Figure 5:**
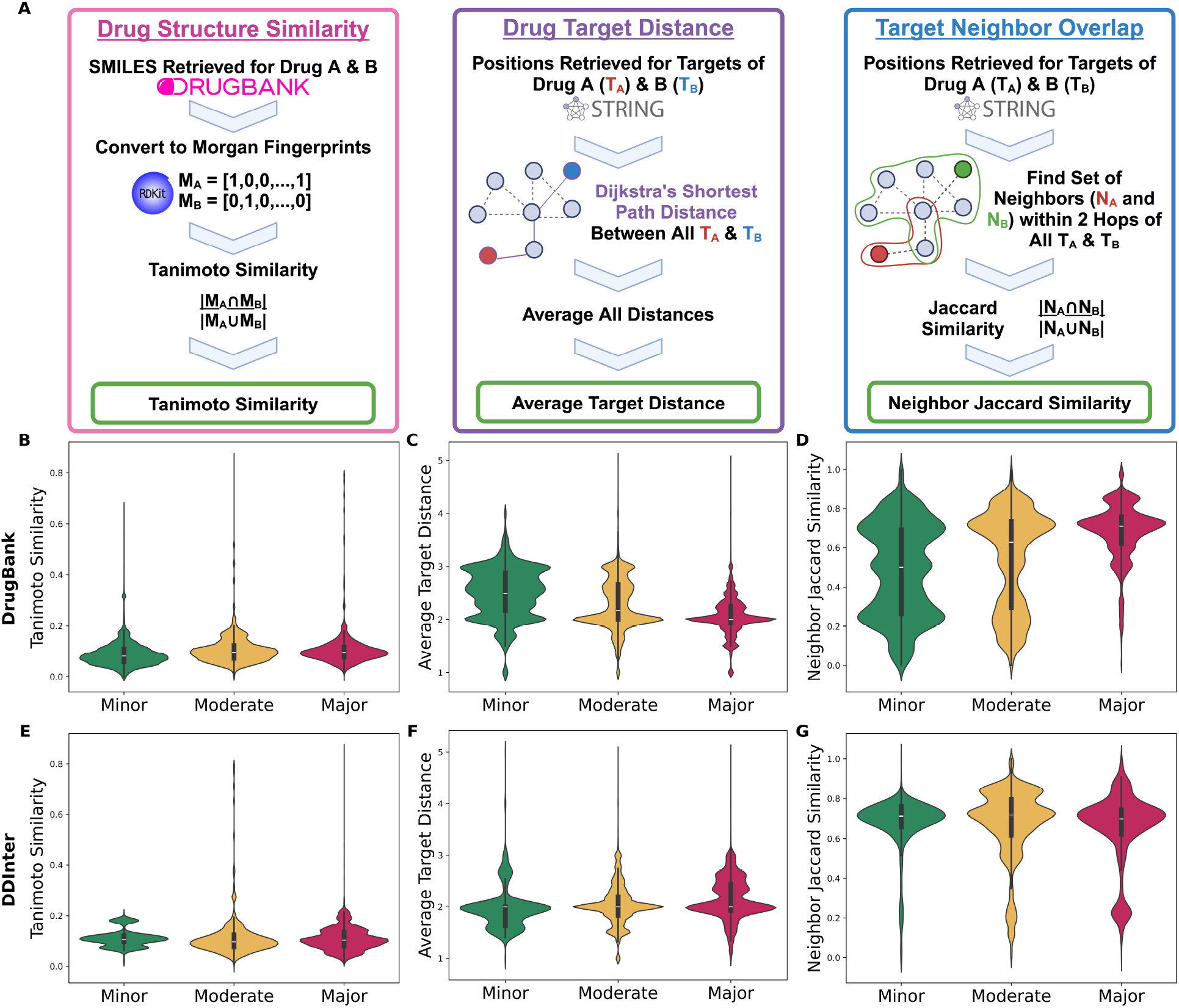
Toxicity metric distributions differ between toxicity categories, but still show large overlaps. Panel **(A)** contains a schematic illustrating how each toxicity metric is calculated. Panels **(B-G)** include violin plots displaying the distributions of three toxicity scoring metrics, split by toxicity categories determined by DrugBank or DDInter, all of which have considerable overlap. The mean of each distribution is denoted by the black horizontal line. All distributions representing the Minor toxicity are in green, the Moderate in yellow, and the Major in red. The first row **(B)**-**(D)** corresponds to toxicity categories determined via the DrugBank database. The second row **(E)**-**(G)**) corresponds to the toxicity categories determined by the DDInter database. The first column (panels **(B)** and **(E)**) shows the Tanimoto Similarity distributions between drug structure Morgan Fingerprint representations in each drug pair. The second column (panels **(C)** and **(F)**) uses the average target distance between the sets of drug targets in a drug combination when Dijkstra’s shortest path is applied to the target locations on the STRING PPIN network. Panels **(D)** and **(G)**, in the last column, displays the distribution of the Jaccard Similarity of the sets of drug targets’ neighbors, within two-hops on the STRING network.

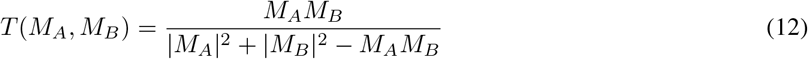

Once all Tanimoto Similarities had been calculated, we plotted the distributions of Morgan Fingerprint similarities in violin plots by toxicity categories from both DrugBank and DDInter.

The second metric we examined was the average distance between targets of each drug in a combination. Since toxicity is thought to be due to the overlap of drug target pathways, we investigated whether larger overall distances between drug targets meant lower severity of DDIs. To compute this average target distance score, we used the STRING PPIN database [41], limited to known interactions (rather than predicted) in *Homo sapiens*. We used the target UniProt IDs retrieved from DrugBank and converted these to STRING IDs using the UniProt database [65]. To find the average target distance for a given drug combination (drug *A* and drug *B*), let *T*_*A*_ = {*t*_*A*,1_, *t*_*A*,2_, …, *t*_*A*,*n*_} as the set of targets for drug *A* and *T*_*B*_ = {*t*_*B*,1_, *t*_*B*,2_, …, *t*_*B*,*m*_} as the set of targets for drug *B*. Next, let *G* = (*V, E*) represent the STRING PPIN where *V* is the set of proteins and *E* is the set of interactions. Let *d*_*G*_(*u, v*) represent Dijkstra’s [66] shortest path distance between the *u*-th and *v*-th proteins. The average target distance between all respective targets of drugs *A* and *B* can be formulated as:

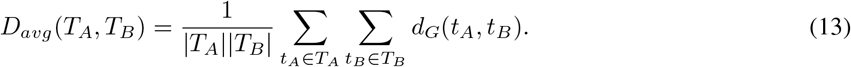

After computing the average target distance for all drug combinations, we then separated the distance values by either DrugBank or DDInter toxicity categories. We again use violin plots to show the distributions of average target distance between minor, moderate, and major toxicity.

The last toxicity score aggregated drug target neighborhood information via the STRING PPIN [15, 35]. Yang et al. [35] used neighborhoods that extended two hops from the original drug target. We assessed whether the overlap of two hop neighborhoods was correlated with known toxicity. For a drug combination (drug *A* and drug *B*) and their set of drug targets (*T*_*A*_ and *T*_*B*_), we found the positions of all targets on the STRING graph *G* = (*V, E*). For each target *t*_*A*,*i*_ ∈ *T*_*A*_, we found all *v* ∈ *V*, where *v* is within two degrees of *t*_*A*,*i*_, creating a neighborhood set of proteins *N*_*A*_. We also found all *v* ∈ *V* that are neighbors for the targets of drug *B*, creating *N*_*B*_. To calculate the neighborhood overlap, we then used the Jaccard Similarity of *N*_*A*_ and *N*_*B*_. As with the other metrics, we separated neighbor Jaccard Similarity by toxicity categories via DrugBank or DDInter and created violin plots of the distributions.

For each of these metrics, we tested for normality via scipy’s `normaltest` function and determined the distributions were not normal (*p* ≤ 0.05). Thus, we used the non-parametric tests of the Kruskal-Wallis and Dunn’s with Bonferroni correction to assess whether different toxicity categories had different toxicity scores medians. We then used the Jonckheere-Terpstra test to assess if toxicity score medians trended with higher or lower toxicity.

## 3 Results

### 3.1 Optimizing for synergy scores alone may prioritize toxic combinations

We investigated whether or not there is any relationship between synergy scores and toxicity levels. This allows us to understand if current methods that optimize for synergy scores alone are independent of toxicity or if higher synergy indicates reduced toxicity. First, we explored the percentages of synergistic combinations belonging to each severity level of DDIs (Figure 2A). It is notable that for all synergy scores, the majority of drug combinations in the DrugBank-DrugComb dataset belong to major adverse DDIs. DrugComb is one of the most commonly used datasets for predicting cancer drug synergy, and if the dataset is biased too heavily to high toxicity combinations, prediction models may be prioritizing more toxic candidates.

Furthermore, using the Kruskal-Wallis test, we found that all synergy scores in the DrugBank dataset—outside of Loewe (*p* = 5.06 × 10^−2^)—had significant differences in their medians between toxicity categories: Bliss (*p* = 5.60 × 10^−168^), HSA (*p* = 2.16 × 10^−77^), ZIP (*p* = 2.28 × 10^−217^), S_max (*p* = 2.24 × 10^−148^), S_mean (*p* = 6.71 × 10^−223^), and S_sum (*p* = 1.25 × 10^−170^). An example distribution of S_mean is shown in Figure 2B, with additional boxplots present in Supplementary Figure 1. This indicated that synergy scores are generally not independent of toxicity. We also computed Dunn’s test (with Bonferroni correction) on each of the synergy score distributions to determine if there were significant differences between the medians of each pair of toxicity categories. In other words, we explored if there differences in a synergy score’s major toxicity distribution compared to its moderate toxicity distribution (major versus minor or moderate versus minor). The remaining synergy scores (Bliss, HSA, ZIP, S_max, S_mean, and S_sum) all consistently found significant differences between each of the pairwise toxicity categories (Supplementary Table 1).

Next, we applied the Jonckheere-Terpstra test to assess whether increasing synergy scores trend with lower toxicity. In the DrugBank database, we observed the opposite. Bliss, HSA, ZIP, S_max, S_mean, and S_sum were all found to have increasing toxicity categories significantly associated with higher synergy scores (*p* ≈ 0.00; Supplementary Table 1). This indicates that current methods prioritizing highly synergistic combinations may actually increase the chances for major DDIs. Notably, the Loewe synergy score did not have any significant difference in the medians of its toxicity distributions, indicating that the relationship between synergy scores and toxicity depends on how synergy is calculated.

However, these observations are not consistent between toxicity databases. The toxicity categories represent very different fractions in the DDInter database, where most synergistic combinations are moderate (Figure 2C). Still, major toxicity represents near a fifth of synergistic combinations across all synergy scores. When the Kruskal-Wallis test is applied to the DDInter toxicity categories (Supplementary Figure 2), HSA (*p* = 1.29 × 10^−113^), S_max (*p* = 2.29 × 10^−73^), S_mean (*p* = 1.52 × 10^−131^), and S_sum (*p* = 3.33 × 10^−22^) agreed with the DrugBank findings where there were significant differences between the synergy score distributions when divided by toxicity category. However, three of the synergy scores had differing conclusions. Bliss (*p* = 0.106) and ZIP (*p* = 0.506) did not have statistically significant differences in DDInter toxicity category distributions while Loewe (*p* = 2.55 × 10^−17^) did. These inconsistencies also appeared within Dunn’s post-hoc tests: the S_mean (Figure 2D) and S_sum did not consistently find significant differences in the distributions when comparing toxicity categories pairwise. Furthermore, via the Jonckheere-Terpstra tests, the HSA and S_max synergy scores’ lower toxicity categories were significantly associated with higher synergy scores—another reversal from DrugBank (Supplementary Figure 2). There was no single synergy score in which the findings from DrugBank and DDInter agreed, indicating that the specific toxicity database drastically affects the synergy score and toxicity relationship. The relationship between synergy and toxicity appears to be score dependent, but our analyses show that prioritizing synergy alone may not lead to optimally low toxicity combinations.

### 3.2 Overlap in drug targets correlates with toxicity but fails to distinguish between toxicity categories

To better understand the biological mechanisms contributing to different DDI severity levels and synergy, we also retrieved drug target and pathway information. Here, we used the Jaccard Similarity to quantify the overlap between a drug combination’s drug targets, pathways, and lowest-level pathways (Figure 3A). We first used the Kruskal-Wallis test to verify if the medians of the Jaccard Similarity distributions differed when split by toxicity category, which we found to be significant (*p <* 0.05) in all similarity classes and both DrugBank and DDInter (Supplementary Table 2). We then assessed whether there were pairwise differences in similarity distributions using Dunn’s test with Bonferroni correction, and we found all post-hoc analyses to also be significant (*p <* 0.05) (Supplementary Table 2). Previous studies have attributed toxicity to high overlap of drug target pathways [15, 35, 36], so our initial hypothesis was that higher Jaccard Similarity in all classes (drug targets, all Reactome pathways, or Reactome’s lowest pathways) would trend with increasing toxicity levels. To test this, we applied the Jonckheere-Terpstra test and found that the median Jaccard Similarities in all classes were significantly associated with higher toxicity levels (*p <* 0.05) (Supplementary Table 2). These findings confirm that overlapping drug targets and their pathways can recapitulate toxicity information in known DDI databases like DrugBank and DDInter.

Because the Jonckheere-Terpstra results indicate that the medians of each overlap distribution increase from minor toxicity to major toxicity, one might consider using the Jaccard Similarity metrics as a cutoff to distinguish drug combinations’ toxicity. However, our strip plot distributions show considerable overlap for each of the Jaccard Similarity classes (Figure 3B-3G). The distributions do not always show that the major toxicity category has the most samples with the highest Jaccard Similarity. In fact, there are combinations in all panels that have complete overlap (Jaccard Similarity equal to 1.0) that exist in the minor category. This indicates that while higher overlap trends with higher toxicity, an overlap metric like Jaccard Similarity does not separate toxicity categories well.

### 3.3 Synergy scores do not correlate with overlap of drug targets and their pathways

In the second aspect of our mechanistic analysis, we also sought to understand whether overlapping drug targets and pathways were correlated with synergy. If certain synergy scores were also associated with drug target overlap, this could account for why some synergy scores trended with higher toxicity. Figure 4 displays scatter plots for DrugBank with the Bliss synergy score on the y-axis and the Jaccard Similarities on the x-axis. The other six synergy scores using the DrugBank dataset were plotted in Supplementary Figure 3. Results for DDInter can be found in Supplementary Figure 4. Interestingly, none of the results show any discernible correlation, with all *R*^2^ values near 0. This observation is confirmed when calculating the Pearson and Spearman correlations, which are compiled in Supplementary Table 3. All Pearson and Spearman correlation coefficients are near zero, indicating no correlation between synergy scores and drug targets or pathways. This analysis demonstrates that drug target overlap alone is not responsible for the trends we observed among certain synergy scores and toxicity.

### 3.4 Toxicity scoring methods struggle to clearly delineate DDI severity

Finally, we assessed common methods for creating toxicity scores, which have been used to create penalty terms in the loss functions of cancer drug combination prediction models. We computed three metrics: the Tanimoto Similarity of Morgan Fingerprint bit vectors, the average target distance, and the neighbor Jaccard Similarity (Figure 5). We also performed the Kruskal-Wallis test with Dunn post-hoc analysis (Bonferroni corrected) on each of the toxicity scores (test statistics and p-values are in Supplementary Table 4).

#### Morgan Fingerprint Tanimoto Similarity

We first established through the Kruskal-Wallis test that each score did have significantly different medians between toxicity categories, regardless of which database was used. Morgan Fingerprint Tanimoto Similarity measures how similar the drug structures are in a drug pair. Dunn post-hoc analysis identified significant differences in each of the pairwise DDInter toxicity categories (Dunn’s Test on Major versus Minor DDInter toxicity *p* = 1.52 × 10^−3^, Major to Moderate toxicity *p* = 1.77 × 10^−20^, and Moderate-Minor *p* = 1.0753 × 10^−13^). However, not all pairwise categories were significant in the DrugBank toxicity data: the medians of Major and Moderate DrugBank toxicity were not found to be significantly different (*p* = 0.0532), although Major to Minor (*p* = 3.72 × 10^−216^) and Moderate to Minor (*p* = 1.45 × 10^−208^) DrugBank toxicity distributions did differ significantly. When the Jonckheere-Terpstra test was applied, both DrugBank and DDInter found a significant correlation of increasing Morgan Fingerprint Tanimoto Similarity with increasing toxicity (DrugBank *p* ≈ 0.00, DDInter *p* = 2.62 × 10^−9^). These findings agree with the hypothesis that higher drug structure similarity trends with a higher chance of severe DDIs. However, the Tanimoto Similarity distributions have a large amount of overlap between the toxicity categories in both DrugBank and DDInter (Figure 5B and 5E). The vast majority of drug combinations also appear to have Tanimoto Similarities between the range of 0 and 0.2.

#### Average Target Distance

We next tested whether the closeness of drug targets on a PPIN could correlate well with toxicity levels. We expected that a high average target distance would be correlated with less severe DDIs. We found that, while there was the DrugBank and DDInter datasets agreed on were significant differences in the medians of the average target distance distributions between toxicity categories (Kruskal-Wallis and Dunn Test results in Supplementary Table 4), there was disagreement when we applied the Jonckheere-Terpstra test. In DrugBank, high target distance significantly trended with lower toxicity (*p* ≈ 0.00), but in DDInter, we found the opposite (*p* ≈ 0.00). Average target distance as a metric is very dependent on complete knowledge of drug targets and their interactions in a PPIN, so it may be more sensitive to a smaller dataset like DDInter and incomplete target information. Furthermore, our results also show overlapping distributions between toxicity categories (e.g., Figure 5C and 5F).

#### Two-hop Neighborhood Jaccard Similarity

Finally, to assess a drug target-based toxicity score that was less dependent on perfect knowledge of drug targets and their interactions, we considered a two-hop neighborhood Jaccard Similarity analysis. We adopted this measure from the toxicity score used by Yang et al. [35], which aggregated neighbors within two degrees of drug targets. Compared to metrics like the average target distance or drug target overlap, the two-hop neighborhood widened the drug target area from a few specific proteins to a small subnetwork within a PPIN. Assessing the overlap of these subnetworks could protect against the potential pitfalls where incomplete target or interaction information may skew the other metrics. In the neighborhood Jaccard Similarity, we found significant differences in the medians among all toxicity categories (DrugBank *p* ≈ 0.00 and DDInter *p* = 7.62 × 10^−23^) for both DDI databases. However, there was disagreement (1) in the neighborhood similarity distributions between Major toxicity and Minor toxicity (DrugBank *p* ≈ 0.00 versus DDInter *p* = 0.83) and (2) in whether higher neighborhood similarity was associated with higher toxicity (DrugBank Jonckheere-Terpstra Increasing Toxicity *p* ≈ 0.00) or lower toxicity (DDInter Jonckheere-Terpstra Decreasing Toxicity *p* = 2.22 × 10^−16^). Furthermore, our results again show a considerable amount of overlap between distributions (e.g., Figure 5D and 5G).

## 4 Discussion

In this study, we explored whether current metrics of synergy and toxicity are well correlated with clinical toxicity in cancer drug combinations, addressing a critical validation need in current computational approaches to combination therapy development. Our findings reveal important limitations in how toxicity is currently incorporated into synergy prediction models. While the field has largely focused on optimizing synergy scores as a primary metric for identifying promising drug combinations, our analysis suggests this may overlook crucial toxicity considerations that impact clinical viability. The complexity of drug-drug interactions extends beyond what current target overlap and structural similarity metrics can capture, highlighting the need for more comprehensive and validated toxicity measurements. These results emphasize the importance of balancing efficacy with safety in a more nuanced and data-driven manner.

First, when examining the DrugComb database, we found a substantial portion of synergistic combinations associated with major adverse drug-drug interactions (DDIs)—even in the most conservative toxicity classification from DDInter, approximately 20% of synergistic combinations showed high toxicity. Our statistical analysis confirmed that certain synergy scores significantly correlate with increased toxicity levels, showing that synergy and toxicity are not strictly independent. Notably, this relationship varies depending on which synergy scoring method is used, highlighting the critical importance of carefully selecting appropriate synergy metrics when evaluating potential combination therapies. These findings suggest an important caution: optimization strategies focused solely on synergy may inadvertently select combinations with higher likelihood of dangerous drug interactions.

Our mechanistic investigation into drug-drug interactions revealed nuanced relationships between molecular features, toxicity, and synergy. While we found minimal correlation between synergy scores and either drug target or pathway overlap, we did confirm significant differences in Jaccard Similarities across toxicity categories. The Jonckheere-Terpstra tests verified a clear trend: higher target and pathway overlap consistently correlated with increased toxicity severity across both DrugBank and DDInter databases. This supports existing understanding that shared targets increase toxic interaction risks. However, our distribution analysis revealed important limitations: some combinations with complete target overlap still exhibited only minor toxicity, demonstrating that overlap metrics alone cannot definitively predict toxicity outcomes. Taken together, our findings suggest that while target and pathway overlap metrics provide valuable insights for toxicity assessment, they appear to have limited utility as predictors of drug synergy.

We evaluated three approaches to toxicity penalties and identified significant limitations with each method. Morgan Fingerprint Tanimoto Similarity consistently correlated with toxicity across both databases, with higher structural similarity trending toward greater toxicity severity. However, its practical utility as a standalone predictor is limited by substantial overlap in distributions between toxicity categories and the narrow range of observed values (mostly 0-0.2). Our analysis of average target distance revealed concerning inconsistencies between DrugBank and DDInter, highlighting database-dependent variability. Similarly, two-hop neighborhood Jaccard Similarity showed significant distributional differences across toxicity categories but with contradictory directional trends between databases. These inconsistencies highlight a fundamental challenge in developing reliable toxicity penalties for synergy prediction models. While these metrics capture some biologically relevant information, none provides sufficiently robust discrimination between toxicity categories to function effectively as a standalone penalty in computational models. This emphasizes the need for integrative approaches that combine multiple toxicity indicators rather than relying on any single structural or network-based metric.

This study has several limitations, the most striking of which is the dependence on known DDI datasets. Throughout the first and third components of our analyses, we often found conflicting determinations depending on the DDI database used. In the analysis examining synergy scores and toxicity categories, no single synergy metric maintained fully consistent relationships across both databases. Additionally, the drug target-based toxicity scores (average target distance and neighborhood Jaccard Similarity) concluded opposite trends depending on whether DrugBank or DDInter was used. These contradictory findings emphasize that these results are highly dependent on the specific toxicity database employed. This also indicates a broader need for agreement between known DDI databases, which is especially important for machine learning methods that make predictions while leveraging prior knowledge. While one source of disparity may be due DrugBank’s more up-to-date curation and large difference in sheer size, future research ought to interrogate the divergence between the two datasets to provide researchers a better understanding of the advantages or biases of using one database over another. Additionally, the availability of high-throughput drug combination toxicity screens on healthy control cells could allow future methods to ground their toxicity penalties in real experimental data.

In conclusion, our study highlights a critical gap in current approaches to cancer drug combination prediction: the inadequate integration of toxicity considerations in synergy-based models. Our findings demonstrate that synergy scores alone may inadvertently bias selection toward more toxic combinations, while common toxicity metrics show inconsistent correlation with clinical adverse effects across different databases. These results underscore the complexity of drug-drug interactions and the limitations of current computational approaches that rely on simplified molecular features or network metrics. Moving forward, we advocate for the development of more comprehensive models that explicitly balance efficacy with safety, incorporating multiple complementary toxicity indicators rather than single metrics. Additionally, we emphasize the urgent need for standardized, high-quality combinatorial toxicity data to support these efforts. Ultimately, we hope this work provides a foundation for more clinically relevant computational methods that can accelerate the discovery of effective cancer combination therapies while prioritizing patient safety.

## 5 Acknowledgments

We thank Julian Stamp (Brown University) for his helpful comments in writing this manuscript. We are also grateful to Professors Barbara Engelhardt (Stanford University), Sean Lawler (Brown University), and Ying Ma (Brown University) for helpful discussions and feedback that guided the early directions of this study. This research was conducted in part using computational resources and services at the Center for Computation and Visualization at Brown University.

## 6 Data and code availability

The DrugComb database is publicly available to download at https://drugcomb.fimm.fi/. Version 1.5 was used for this study. For toxicity data, we downloaded the DrugBank (version 5.1.12) including the clinical data under an academic license, which can be found at https://go.drugbank.com/. We also used DDInter, where we downloaded all files in April 2024 from https://ddinter.scbdd.com/download/. Reactome was accessed in December 2024 to map UniProt IDs to human pathways both in the lowest level of pathway as well as all pathways from https://reactome.org/download-data. The STRING PPIN database at https://stringdb-downloads.org/download/protein.physical.links.detailed.v12.0/9606.protein.physical.links.detailed.v12.0.txt.gz was limited to the known interactions in *Homo sapiens*, using version 12.0.

The code written for this project is available at https://github.com/amw14/toxicity-cancer-drug-combination. The RDKit package used to retrieve the Morgan Fingerprint representations of all drug structures was installed using the 2024.09.05 documentation (https://www.rdkit.org/docs/GettingStartedInPython.html).

## 8 Author contributions

AMW and LC conceived the study. AMW wrote the code and performed the analysis. LC supervised the project and provided resources. All authors wrote and revised the manuscript.

