## Supplement for "Characterizing Clinical Toxicity in Cancer Combination Therapies"

A Preprint

**Alexandra M. Wong**
Brown University
Providence, RI 02912


**Lorin Crawford**
Microsoft Research
Cambridge, MA 02142


### 1 Supplementary Material

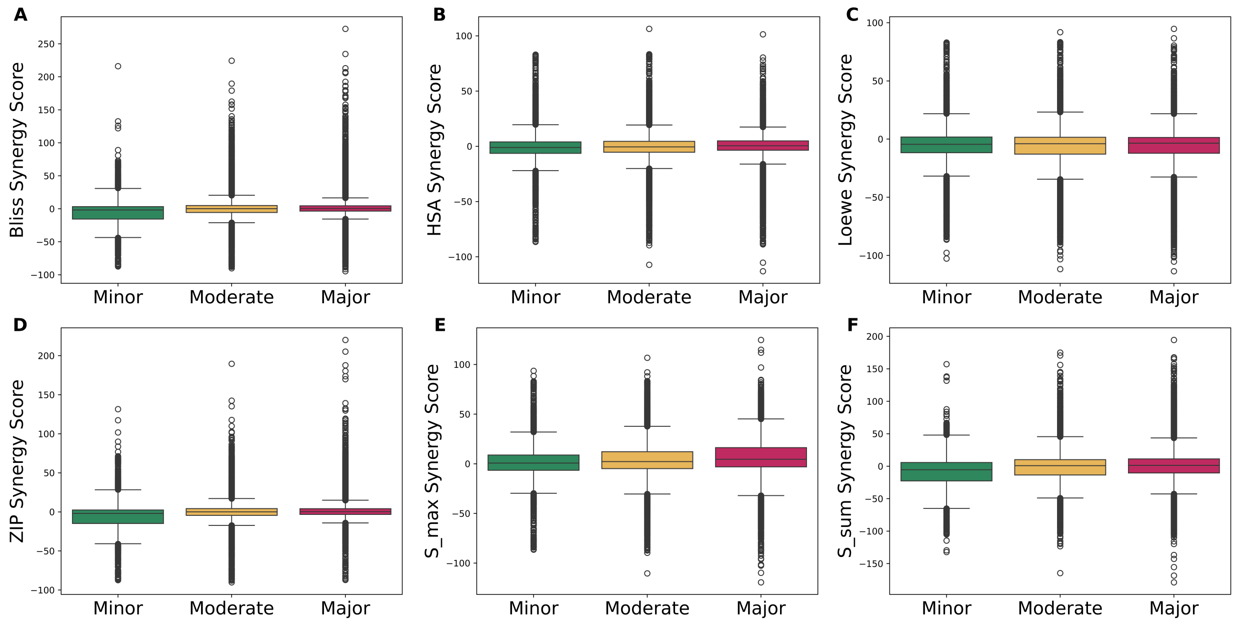


**Supplementary Figure 1: Remaining synergy score distributions show overlap between DrugBank toxicity categories.** Strip plots showing the distributions of drug combination synergy scores when split across toxicity categories, the mean of each distribution is denoted by the black horizontal line. All distributions representing the Minor toxicity are in green, the Moderate in yellow, and the Major in red. Each panel represents a different synergy score: **(A)** the Loewe synergy score, **(B)** S_max, and **(C)** S_sum.

**Supplementary Table 1: Statistical analysis of synergy scores by toxicity category.** Results of statistical analysis assessing the relationship between synergy score distributions and known DDI severity categories. Each section corresponds to a different synergy scoring method: Bliss, HSA, Loewe, ZIP, S_max, S_mean, or S_sum. The results for the Kruskal-Wallis, Dunn with Bonferroni correction, and Jonckheere-Terpstra tests are included for both the DrugBank and DDInter datasets. The Jonckheere-Terpestra Increasing Toxicity test assesses whether the distribution of the Jaccard Similarity increases when toxicity categories become more severe, while the Jonckheere-Terpstra Decreasing Toxicity test evaluates if the Jaccard Similarity distributions increase when the toxicity categories decrease in severity.

| Synergy Score | Test | DrugBank | | DDInter | |
|---|---|---|---|---|---|
| | | Test Statistic | P-value | Test Statistic | P-value |
| Bliss | Kruskal-Wallis | 7.7022e+02 | 5.6027e-168 | 4.4914e+00 | 1.0585e-01 |
| | Dunn: Major/Minor | - | 8.0827e-168 | - | 1.0887e-01 |
| | Dunn: Major/Moderate | - | 1.2205e-17 | - | 9.9011e-01 |
| | Dunn: Moderate/Minor | - | 1.0490e-91 | - | 2.1936e-01 |
| | Jonckheere-Terpstra: Increasing Toxicity | 2.2832e+01 | 0.0000e+00 | 1.6444e+00 | 5.0042e-02 |
| | Jonckheere-Terpstra: Decreasing Toxicity | -2.2832e+01 | 1.0000e+00 | -1.6444e+00 | 9.4996e-01 |
| HSA | Kruskal-Wallis | 3.5306e+02 | 2.1570e-77 | 5.1988e+02 | 1.2855e-113 |
| | Dunn: Major/Minor | - | 2.8117e-50 | - | 2.1276e-02 |
| | Dunn: Major/Moderate | - | 4.5996e-47 | - | 4.6788e-94 |
| | Dunn: Moderate/Minor | - | 3.5059e-06 | - | 7.6845e-30 |
| | Jonckheere-Terpstra: Increasing Toxicity | 1.8743e+01 | 0.0000e+00 | -1.5109e+01 | 1.0000e+00 |
| | Jonckheere-Terpstra: Decreasing Toxicity | -1.8743e+01 | 1.0000e+00 | 1.5109e+01 | 0.0000e+00 |
| Loewe | Kruskal-Wallis | 5.9671e+00 | 5.0613e-02 | 7.6413e+01 | 2.5539e-17 |
| | Dunn: Major/Minor | - | 4.2288e-01 | - | 2.9690e-05 |
| | Dunn: Major/Moderate | - | 7.6071e-02 | - | 2.5699e-08 |
| | Dunn: Moderate/Minor | - | 1.0000e+00 | - | 5.5296e-12 |
| | Jonckheere-Terpstra: Increasing Toxicity | 2.3468e+00 | 9.4686e-03 | -2.6866e+00 | 9.9639e-01 |
| | Jonckheere-Terpstra: Decreasing Toxicity | -2.3468e+00 | 9.9053e-01 | 2.6866e+00 | 3.6087e-03 |
| ZIP | Kruskal-Wallis | 9.9768e+02 | 2.2784e-217 | 1.3605e+00 | 5.0649e-01 |
| | Dunn: Major/Minor | - | 1.1561e-217 | - | 9.0038e-01 |
| | Dunn: Major/Moderate | - | 1.5923e-19 | - | 1.0000e+00 |
| | Dunn: Moderate/Minor | - | 1.6883e-123 | - | 1.0000e+00 |
| | Jonckheere-Terpstra: Increasing Toxicity | 2.5577e+01 | 0.0000e+00 | -1.0900e+00 | 8.6215e-01 |
| | Jonckheere-Terpstra: Decreasing Toxicity | -2.5577e+01 | 1.0000e+00 | 1.0900e+00 | 1.3785e-01 |
| S_max | Kruskal-Wallis | 6.7995e+02 | 2.2418e-148 | 3.3452e+02 | 2.2926e-73 |
| | Dunn: Major/Minor | - | 1.6916e-106 | - | 2.2169e-08 |
| | Dunn: Major/Moderate | - | 8.7756e-78 | - | 5.2444e-48 |
| | Dunn: Moderate/Minor | - | 9.0896e-18 | - | 1.1429e-33 |
| | Jonckheere-Terpstra: Increasing Toxicity | 2.5959e+01 | 0.0000e+00 | -9.1327e+00 | 1.0000e+00 |
| | Jonckheere-Terpstra: Decreasing Toxicity | -2.5959e+01 | 1.0000e+00 | 9.1327e+00 | 0.0000e+00 |
| S_mean | Kruskal-Wallis | 1.0231e+03 | 6.7127e-223 | 6.0244e+02 | 1.5232e-131 |
| | Dunn: Major/Minor | - | 1.5347e-146 | - | 3.1849e-01 |
| | Dunn: Major/Moderate | - | 3.8066e-130 | - | 7.9790e-114 |
| | Dunn: Moderate/Minor | - | 4.8272e-18 | - | 1.4193e-28 |
| | Jonckheere-Terpstra: Increasing Toxicity | 3.2064e+01 | 0.0000e+00 | -1.7241e+01 | 1.0000e+00 |
| | Jonckheere-Terpstra: Decreasing Toxicity | -3.2064e+01 | 1.0000e+00 | 1.7241e+01 | 0.0000e+00 |
| S_sum | Kruskal-Wallis | 7.8244e+02 | 1.2470e-170 | 9.8907e+01 | 3.3320e-22 |
| | Dunn: Major/Minor | - | 1.1982e-171 | - | 7.1489e-06 |
| | Dunn: Major/Moderate | - | 1.4369e-11 | - | 3.4563e-22 |
| | Dunn: Moderate/Minor | - | 1.2305e-104 | - | 1.0000e+00 |
| | Jonckheere-Terpstra: Increasing Toxicity | 2.2079e+01 | 0.0000e+00 | 9.6399e+00 | 0.0000e+00 |
| | Jonckheere-Terpstra: Decreasing Toxicity | -2.2079e+01 | 1.0000e+00 | -9.6399e+00 | 1.0000e+00 |

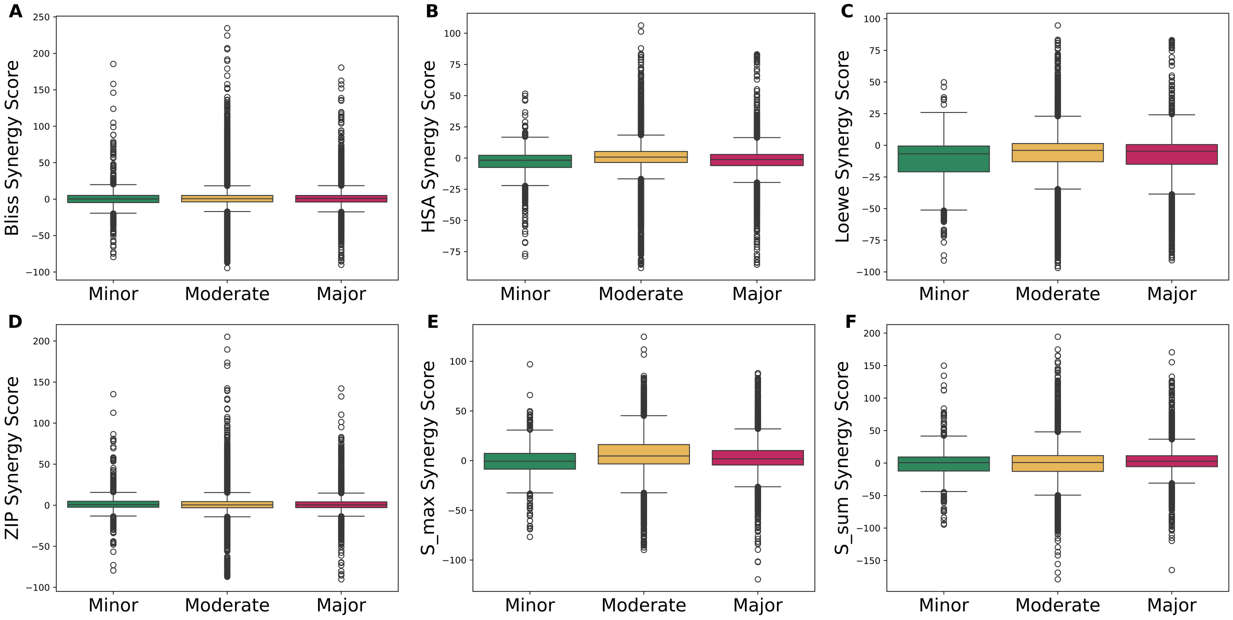


**Supplementary Figure 2: Synergy score distributions by DDInter toxicity category.** Strip plots showing the distributions of drug combination synergy scores when split across toxicity categories, the mean of each distribution is denoted by the black horizontal line. All distributions representing the Minor toxicity are in green, the Moderate in yellow, and the Major in red. Each panel represents a different synergy score: **(A)** the Bliss synergy score, **(B)** HSA, **(C)** Loewe, **(D)** ZIP, **(E)** S_max, **(F)** S_mean, and **(G)** S_sum.

**Supplementary Table 2: Drug target and pathway overlap toxicity analysis**. Results for statistical analysis of drug targets and pathway overlap correlated with toxicity categories. Each section refers to a different Jaccard Similarity metric. The first section computed the overlap between the sets of drug targets in a given drug combination, the second the sets of pathways, and the third the sets of pathways when restricted to the lowest level of Reactome. The results for the Kruskal-Wallis, Dunn with Bonferroni correction, and Jonckheere-Terpstra tests are included for both the DrugBank and DDInter datasets. The Jonckheere-Terpestra Increasing Toxicity test assesses whether the distribution of the Jaccard Similarity increases when toxicity categories become more severe, while the Jonckheere-Terpstra Decreasing Toxicity test evaluates if the Jaccard Similarity distributions increase when the toxicity categories decrease in severity.

| **Overlap Metric** | **Test** | **DrugBank** | | **DDInter** | |
|---|---|---|---|---|---|
| | | **Test Statistic** | **P-value** | **Test Statistic** | **P-value** |
| **Drug Target Jaccard Similarity** | | | | | |
| Drug Target Jaccard | Kruskal-Wallis | 1.5326e+02 | 5.2536e-34 | 1.5647e+02 | 1.0525e-34 |
| | Dunn: Major/Minor | - | 9.4915e-31 | - | 1.3521e-25 |
| | Dunn: Major/Moderate | - | 2.6362e-11 | - | 9.0799e-22 |
| | Dunn: Moderate/Minor | - | 1.6838e-10 | - | 5.7192e-12 |
| | Jonckheere-Terpstra: Increasing Toxicity | 4.7818e+00 | 8.6869e-07 | 5.6787e+00 | 6.7866e-09 |
| | Jonckheere-Terpstra: Decreasing Toxicity | -4.7818e+00 | 1.0000e+00 | -5.6787e+00 | 1.0000e+00 |
| **All Reactome Pathways Jaccard Similarity** | | | | | |
| All Reactome Pathways Jaccard | Kruskal-Wallis | 1.0964e+03 | 8.2813e-239 | 7.6788e+02 | 1.8097e-167 |
| | Dunn: Major/Minor | - | 1.0391e-191 | - | 1.0000e+00 |
| | Dunn: Major/Moderate | - | 2.9285e-100 | - | 2.1182e-153 |
| | Dunn: Moderate/Minor | - | 3.3177e-45 | - | 1.2376e-25 |
| | Jonckheere-Terpstra: Increasing Toxicity | 3.2245e+01 | 0.0000e+00 | 2.0789e+01 | 0.0000e+00 |
| | Jonckheere-Terpstra: Decreasing Toxicity | -3.2245e+01 | 1.0000e+00 | -2.0789e+01 | 1.0000e+00 |
| **Lowest Reactome Pathways Jaccard Similarity** | | | | | |
| Lowest Reactome Pathways Jaccard | Kruskal-Wallis | 1.0848e+03 | 2.7898e-236 | 1.5841e+03 | 0.0000e+00 |
| | Dunn: Major/Minor | - | 1.9012e-182 | - | 2.1407e-01 |
| | Dunn: Major/Moderate | - | 6.4413e-108 | - | 0.0000e+00 |
| | Dunn: Moderate/Minor | - | 3.9843e-38 | - | 1.1862e-44 |
| | Jonckheere-Terpstra: Increasing Toxicity | 2.7895e+01 | 0.0000e+00 | 2.7099e+01 | 0.0000e+00 |
| | Jonckheere-Terpstra: Decreasing Toxicity | -2.7895e+01 | 1.0000e+00 | -2.7099e+01 | 1.0000e+00 |

**Supplementary Table 3: Correlation analysis between drug target and pathway Jaccard Similarity and synergy scores.** Results for statistical analysis of drug targets and pathway overlap correlated with synergy scores. Each section refers to a different Jaccard Similarity metric. The first section computed the overlap between the sets of drug targets in a given drug combination, the second the sets of pathways, and the third the sets of pathways when restricted to the lowest level of Reactome. The results for the Kruskal-Wallis, Dunn with Bonferroni correction, and Jonckheere-Terpstra tests are included for both the DrugBank and DDInter datasets. The Jonckheere-Terpestra Increasing Toxicity test assesses whether the distribution of the Jaccard Similarity increases when toxicity categories become more severe, while the Jonckheere-Terpstra Decreasing Toxicity test evaluates if the Jaccard Similarity distributions increase when the toxicity categories decrease in severity.

| **Drug Target Jaccard Similarity** | | | |
|---|---|---|---|
| **Synergy Score** | **Pearson** | **Spearman** | $\mathbf{R^2}$ |
| Bliss | $-1.7625 \times 10^{-2}$ | $-4.0920 \times 10^{-3}$ | $3.1063 \times 10^{-4}$ |
| HSA | $-7.8355 \times 10^{-3}$ | $-2.6949 \times 10^{-2}$ | $6.1396 \times 10^{-5}$ |
| Loewe | $6.3471 \times 10^{-2}$ | $7.1984 \times 10^{-2}$ | $4.0285 \times 10^{-3}$ |
| ZIP | $-1.9607 \times 10^{-2}$ | $-6.4934 \times 10^{-3}$ | $3.8443 \times 10^{-4}$ |
| S_max | $2.0593 \times 10^{-2}$ | $-8.4737 \times 10^{-3}$ | $4.2407 \times 10^{-4}$ |
| S_mean | $-7.8945 \times 10^{-3}$ | $-4.5837 \times 10^{-2}$ | $6.2323 \times 10^{-5}$ |
| S_sum | $2.8955 \times 10^{-3}$ | $1.2554 \times 10^{-2}$ | $8.3836 \times 10^{-6}$ |
| **All Reactome Pathways Jaccard Similarity** | | | |
| **Synergy Score** | **Pearson** | **Spearman** | $\mathbf{R^2}$ |
| Bliss | $-1.4575 \times 10^{-3}$ | $3.4744 \times 10^{-2}$ | $2.1242 \times 10^{-6}$ |
| HSA | $-5.6955 \times 10^{-3}$ | $-2.7964 \times 10^{-2}$ | $3.2439 \times 10^{-5}$ |
| Loewe | $6.8593 \times 10^{-2}$ | $2.9483 \times 10^{-2}$ | $4.7050 \times 10^{-3}$ |
| ZIP | $-2.9755 \times 10^{-4}$ | $3.7445 \times 10^{-2}$ | $8.8535 \times 10^{-8}$ |
| S_max | $2.2987 \times 10^{-2}$ | $1.6085 \times 10^{-2}$ | $5.2840 \times 10^{-4}$ |
| S_mean | $3.2041 \times 10^{-3}$ | $-1.6027 \times 10^{-2}$ | $1.0266 \times 10^{-5}$ |
| S_sum | $2.5985 \times 10^{-2}$ | $8.2550 \times 10^{-2}$ | $6.7524 \times 10^{-4}$ |
| **Lowest Reactome Pathway Jaccard Similarity** | | | |
| **Synergy Score** | **Pearson** | **Spearman** | $\mathbf{R^2}$ |
| Bliss | $-6.3913 \times 10^{-3}$ | $2.8560 \times 10^{-2}$ | $4.0849 \times 10^{-5}$ |
| HSA | $-3.1004 \times 10^{-3}$ | $-4.2875 \times 10^{-2}$ | $9.6123 \times 10^{-6}$ |
| Loewe | $7.5989 \times 10^{-2}$ | $5.7598 \times 10^{-2}$ | $5.7744 \times 10^{-3}$ |
| ZIP | $-6.0386 \times 10^{-3}$ | $2.6652 \times 10^{-2}$ | $3.6465 \times 10^{-5}$ |
| S_max | $1.5437 \times 10^{-2}$ | $-1.1151 \times 10^{-2}$ | $2.3831 \times 10^{-4}$ |
| S_mean | $-9.8367 \times 10^{-5}$ | $-5.2749 \times 10^{-2}$ | $9.6761 \times 10^{-9}$ |
| S_sum | $7.9356 \times 10^{-3}$ | $5.8192 \times 10^{-2}$ | $6.2973 \times 10^{-5}$ |

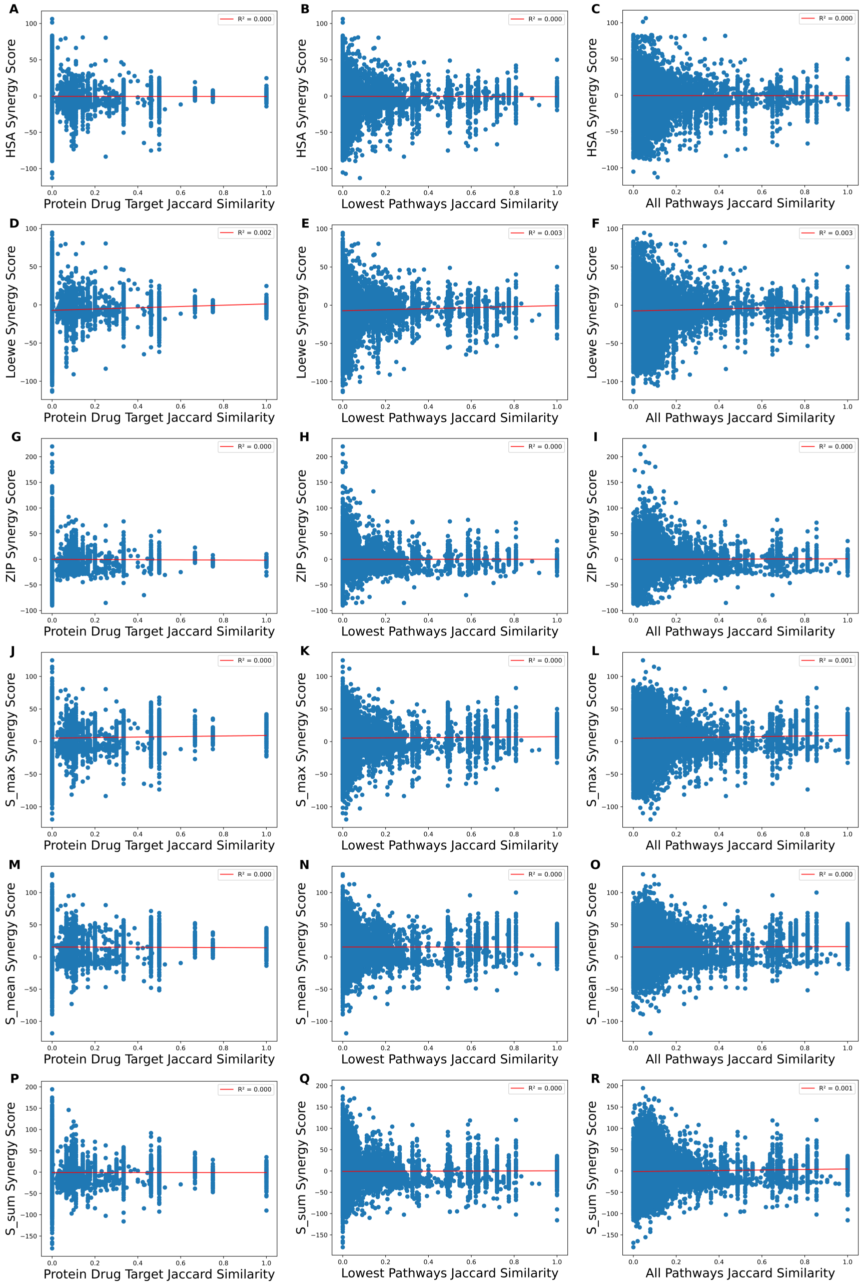


**Supplementary Figure 3: Synergy scores show no correlation with drug target overlap metrics from DrugBank.** Scatter plots showing the relationships between remaining synergy scoring metrics and drug combination target overlaps for drug combinations present in both DrugComb and DrugBank. Each plot also contains a red line for the best fit line, with the $R^2$ present in the legend. Each row corresponds to a different synergy scoring method, and each columns corresponds to a different drug combination target metric. Rows starting with **(A)**, **(D)**, **(G)**, **(J)**, **(M)**, and **(P)** correspond to the HSA, Loewe, ZIP, S_max, S_mean, and S_sum synergy scores. Columns starting with **(A)**, **(B)**, and **(C)** correspond to the Jaccard Similarity of drug combinations' protein targets, pathways when restricted to the lowest level of Reactome, and all pathways of Reactome.

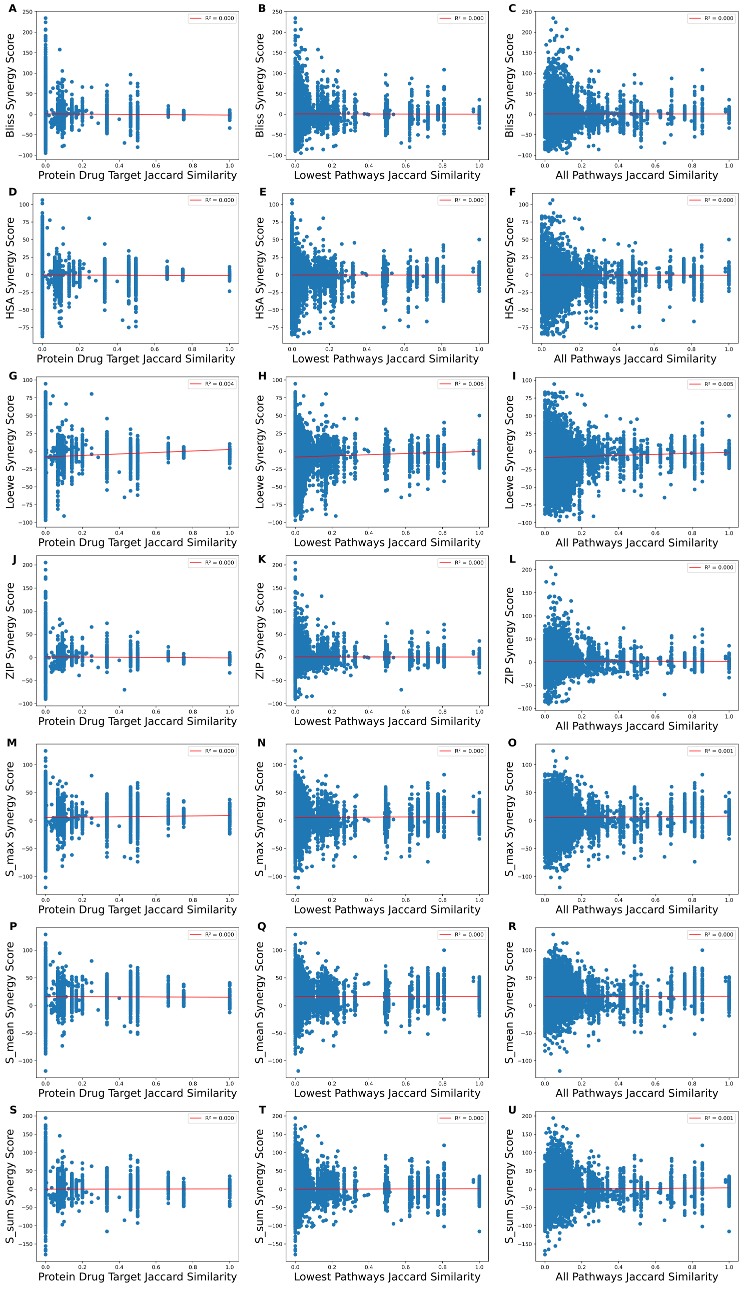


**Supplementary Figure 4: Synergy scores show no correlation with drug target overlap metrics from DDInter.** Scatter plots showing the relationships between synergy scoring metrics and drug combination target overlaps for drug combinations present in both DrugComb and DDInter. Each plot also contains a red line for the best fit line, with the $R^2$ present in the legend. Each row corresponds to a different synergy scoring method, and each columns corresponds to a different drug combination target metric. Rows starting with **(A)**, **(D)**, **(G)**, **(J)**, **(M)**, **(P)**, and **(S)** correspond to the Bliss, HSA, Loewe, ZIP, S_max, S_mean, and S_sum synergy scores. Columns starting with **(A)**, **(B)**, and **(C)** correspond to the Jaccard Similarity of drug combinations' protein targets, pathways when restricted to the lowest level of Reactome, and all pathways of Reactome.

**Supplementary Table 4: Toxicity scoring metrics compared to toxicity categories.** Table containing the results from statistical assessment of whether common principles used in toxicity scores trend with known DDI categories. The first section tests the similarity of drug structure in a combination by calculating the Tanimoto Similarity of the Morgan Fingerprint representations of both drugs in a combination. The second section examines whether closer average target distance in a drug combination is associated with toxicity levels. Finally, the last section calculates the overlap (Jaccard Similarity) of each drug target's neighborhood within two hops of the protein target on the STRING PPIN. The results for the Kruskal-Wallis, Dunn with Bonferroni correction, and Jonckheere-Terpstra tests are included for both the DrugBank and DDInter datasets. The Jonckheere-Terpestra Increasing Toxicity assesses whether the distribution of the toxicity scoring metric increases when toxicity categories become more severe, while the Jonckheere-Terpstra Decreasing Toxicity test evaluates if the toxicity scoring metric distributions increase when the toxicity categories decrease in severity.

| Toxicity Metric | Test | DrugBank | | DDInter | |
|---|---|---|---|---|---|
| | | Test Statistic | P-value | Test Statistic | P-value |
| **Morgan Fingerprint Tanimoto Similarity** | | | | | |
| Morgan Fingerprint Tanimoto | Kruskal-Wallis | 1.0844e+03 | 3.3778e-236 | 1.3442e+02 | 6.4813e-30 |
| | Dunn: Major/Minor | - | 3.7223e-216 | - | 1.5156e-03 |
| | Dunn: Major/Moderate | - | 5.3174e-02 | - | 1.7664e-20 |
| | Dunn: Moderate/Minor | - | 1.4479e-208 | - | 1.0753e-13 |
| | Jonckheere-Terpstra: Increasing Toxicity | 1.7961e+01 | 0.0000e+00 | 5.8393e+00 | 2.6217e-09 |
| | Jonckheere-Terpstra: Decreasing Toxicity | -1.7961e+01 | 1.0000e+00 | -5.8393e+00 | 1.0000e+00 |
| **Average Target Distance** | | | | | |
| Average Target Distance | Kruskal-Wallis | 6.6724e+03 | 0.0000e+00 | 4.6187e+02 | 5.0856e-101 |
| | Dunn: Major/Minor | - | 0.0000e+00 | - | 8.7168e-59 |
| | Dunn: Major/Moderate | - | 0.0000e+00 | - | 6.9037e-76 |
| | Dunn: Moderate/Minor | - | 1.6214e-274 | - | 3.2316e-20 |
| | Jonckheere-Terpstra: Increasing Toxicity | -7.8777e+01 | 1.0000e+00 | 2.1252e+01 | 0.0000e+00 |
| | Jonckheere-Terpstra: Decreasing Toxicity | 7.8777e+01 | 0.0000e+00 | -2.1252e+01 | 1.0000e+00 |
| **Two Hop Neighborhood Jaccard Similarity** | | | | | |
| 2-Hop Neighboring Proteins Jaccard | Kruskal-Wallis | 6.1532e+03 | 0.0000e+00 | 1.0186e+02 | 7.6178e-23 |
| | Dunn: Major/Minor | - | 0.0000e+00 | - | 8.3010e-01 |
| | Dunn: Major/Moderate | - | 0.0000e+00 | - | 1.8138e-22 |
| | Dunn: Moderate/Minor | - | 1.5841e-99 | - | 8.7452e-03 |
| | Jonckheere-Terpstra: Increasing Toxicity | 7.7638e+01 | 0.0000e+00 | -8.1536e+00 | 1.0000e+00 |
| | Jonckheere-Terpstra: Decreasing Toxicity | -7.7638e+01 | 1.0000e+00 | 8.1536e+00 | 2.2204e-16 |
